# *Pannexin 3* deletion in mice results in knee osteoarthritis and intervertebral disc degeneration after forced treadmill running

**DOI:** 10.1101/2023.03.20.532801

**Authors:** Brent Wakefield, Jeffrey Lawrence Hutchinson, Justin Tang, Rehanna Kanji, Courtney Brooks, Cheryle A Séguin, Silvia Penuela, Frank Beier

## Abstract

Pannexin 3 (*Panx3*) is a glycoprotein that forms mechanosensitive channels expressed in chondrocytes and annulus fibrosus cells of the intervertebral disc (IVD). Evidence suggests *Panx3* plays contrasting roles in traumatic versus aging osteoarthritis (OA) and intervertebral disc degeneration (IDD). However, whether its deletion influences the response of joint tissue to mechanical stress is unknown. The purpose of this study was to determine if *Panx3* deletion in mice causes increased knee joint OA and IDD after forced treadmill running. Male and female wildtype (WT) and *Panx3* knockout (KO) mice were randomized to either a no exercise group (sedentary; SED) or daily forced treadmill running (forced exercise; FEX) from 24 to 30 weeks of age. Knee cartilage, tibial secondary ossification center and IVD histopathology were evaluated by histology. Both male and female *Panx3* KO mice developed larger superficial defects of the tibial cartilage after forced treadmill running compared to SED WT mice. Additionally, both male and female *Panx3* KO mice developed greater bone area of the tibial secondary ossification center with running. In the lower lumbar spine, both male and female *Panx3* KO mice developed histopathological features of IDD after running compared to SED WT mice. These findings suggest that the combination of deleting *Panx3* and forced treadmill running induces OA and causes histopathological changes associated with degeneration of the IVDs in mice.

## INTRODUCTION

Knee osteoarthritis (OA) [1] and low back pain [2] are two major causes of disability worldwide. While a subset of these conditions is caused by trauma, they are also associated with genetic [3] and environmental factors (ex. physical activity [4]), often developing spontaneously. Most studies of genetically modified animal models (mice) induce joint stress using injury or surgical interventions, which may not simulate primary OA or spontaneous intervertebral disc degeneration (IDD) in humans. Indeed, the molecular mechanisms of traumatic versus idiopathic joint pathology may differ, highlighting the need to use multiple models of OA or IDD to investigate the role of target genes associated with disease pathogenesis.

Pannexin 3 (PANX3), a membrane bound channel-forming glycoprotein expressed in osteoblasts [5], chondrocytes [6], and annulus fibrosus cells of the IVD [7], has previously been identified as a potential target for OA [8, 9] and IDD [10, 11]. PANX3 expression is induced by the transcription factor RUNX2, which drives chondrocyte hypertrophy [12]. Hypertrophic chondrocytes produce catabolic enzymes such as matrix metalloproteinase (MMP13) which contribute to the breakdown of cartilage extracellular matrix [13]. Our lab has shown that global and chondrocyte specific deletion of *Panx3* in male mice is protective against OA caused by destabilization of the medial meniscus (DMM) surgery (a model of post-traumatic OA) [8].

Paradoxically, we subsequently found that *Panx3* KO mice had worse knee cartilage degradation and sclerotic subchondral bone when aged to 18 months [9]. These data were amongst the first to show completely opposite roles of the same gene in different models of OA – *Panx3* appears to promote OA in a post-traumatic model but protects joints during aging.

With regards to IVD health, *Panx3* KO discs were also protected from trauma-induced degeneration and did not develop spontaneous age-associated IDD [10]. Interestingly, IVDs adjacent to the site of injury experience altered mechanics, and these IVDs of *Panx3* KO mice showed accelerated degeneration compared to wildtype (WT) discs. Taken together, this suggests the absence of PANX3 is beneficial in traumatic models, while its presence is necessary for the adaptive cellular responses to altered or accumulated loading.

In this study we sought to determine how the knee joint and lumbar spine of adult male and female *Panx3* KO mice responded to 6 weeks of daily forced treadmill running. Treadmill running in male WT mice produced superficial cartilage defects to a degree that would be expected in this intervention [14]. While the deletion of *Panx3* in male and female mice seemed to produce cartilage defects, treadmill running had an additive effect that appeared to cause large superficial cartilage defects on average. Additionally, the secondary ossification center of the proximal tibial in *Panx3* KO mice displayed greater bone area after treadmill running compared to sedentary *Panx3* KO mice. In the lumbar spine, forced running in *Panx3* KO mice resulted in more degeneration of IVDs compared to SED WT mice. This evidence suggests an additive effect of treadmill running and *Panx3* deletion on structural damage to knee cartilage and IVDs in mice.

## METHODS

### Mice

All animals used in this study were bred in-house and were raised and euthanized in accordance with the ethics guidelines set forth by the Canadian Council for Animal Care. Animal use protocols were approved by the Council for Animal Care at Western University Canada (AUP 2019-069). Mice were housed in standard shoe box-style caging and exposed to a 12-hour light/dark cycle. Mice ate regular chow ad libitum. Mice were weighed bi-weekly and body composition was assessed at 24 and 30 weeks of age, which was published previously [15]. WT and *Panx3* KO mice were congenic. DNA was collected from ear clippings of each mouse to determine genotype using polymerase chain reaction as previously described [8, 16]. At sacrifice, mouse knees and spines were collected and immediately processed for histological analysis.

### Forced Exercise Intervention

At 24 weeks of age mice were randomized to either a no exercise group (SED) or a forced treadmill running (FEX) group. The FEX groups ran on a treadmill (Columbus Instruments, Ohio) for 6 weeks, 1 hour a day, 5 days a week, at a speed of 11 m/min, and a 10° incline, an adapted protocol that has previously been used to induce osteoarthritis in C57BL6 mice [14]. The mice were encouraged to run using a bottle brush bristle and a shock grid at the end of the treadmill as per the animal ethics protocol.

### Histopathological Assessment of the Knee Joint

Knee joints of the mice were fixed in 4% paraformaldehyde at room temperature for 24 hours on a shaker, and then decalcified in 5% EDTA for 12 days at room temperature. Knees were processed and embedded coronally in paraffin, and 5-μm–thick sections were cut from front to back through the width of the joint. Sections were stained with toluidine blue. Six sections spanning the width of the knee were scored by 2 blinded reviewers using a 12-point system developed and described by McNulty et al. [17]. The average max score of each sample was then used for statistical analysis. Cartilage area of the load bearing region was analysed using the OsteoMeasure (OsteoMetric, Atlanta, GA) software. Unmineralized and mineralized cartilage was manually segmented at the tidemark and analyzed separately.

### Histopathological Assessment of the Lumbar Intervertebral Discs

Lumbar spines harvested for histological analyses were fixed overnight with 4% (w/v) paraformaldehyde, followed by 7 days of decalcification with Shandon’s TBD-2 (Thermo Scientific, Waltham, MA, USA) at room temperature. Following standard processing, tissues were embedded in paraffin and sectioned in the sagittal plane at a thickness of 5 μm. Mid-sagittal sections were deparaffinized and rehydrated as previously described [18] and stained using 0.1% Safranin-O/0.05% Fast Green. Sections were imaged on a Leica DM1000 microscope, with Leica Application Suite (Leica Microsystems: Wetzlar, DEU). To evaluate IVD degeneration, spine sections were scored by two observers blinded to age, exercise, sex, and genotype using a previously established histopathological scoring system for mouse IVDs [19]. Modifications were made in Part (I) (“score 4: mineralized matrix in NP” was omitted). Compartment scores (NP, AF, and NP/AF Boundary) were summed for each IVD and reported for each lumbar level. To report on degeneration across the lumbar spine, scores for individual lumbar IVDs (L2-L6) were summed, and the total score plotted for each individual mouse.

### Measurements of Tibial Subchondral Bone and L5 Vertebral Body

OsteoMeasure (OsteoMetric, Atlanta, GA) software was used to measure the tibial secondary ossification center bone area and the L5 bone area (cortical and trabecular) and marrow area. Three randomly selected slides per animal were chosen, and for the subchondral tibia, the bone tissue was segmented from the overlaying articular cartilage of the tibia and the underlaying growth plate. Bone area and marrow area were divided by the total secondary ossification center area for correction.

### Statistics

The department of Epidemiology and Biostatistics at Western University was consulted to determine the appropriate statistical analysis. Data are presented as means ± CI. Prism (GraphPad Software Inc.) was used to run all statistical tests including one-way analysis of variance (ANOVA) or three-way ANOVA for comparison. For analysis of the articular cartilage structure scores and histopathological scores of the IVD, males and females were analysed separately within their respective genotype and activity group. A Kruskal-Wallis test was used with an uncorrected Dunn’s test for multiple comparisons to determine statistical differences from the SED WT control group. For the cartilage area, subchondral bone, and L5 bone measures, a three-way ANOVA followed by Sidak’s multiple comparisons test was performed to determine the effect of activity within each genotype for each sex. All applicable data met assumptions for homoscedasticity or normality of residuals. Based on the recommendations of the editorial entitled: Moving to a World Beyond “p < 0.05” [20] we did not set a threshold for significance.

## RESULTS

### Forced treadmill running and *Panx3* deletion increases features of knee tibial OA

To better understand how *Panx3* deletion influences OA pathogenesis, we examined the knee joints of male and female WT and *Panx3* KO mice under SED and FEX conditions (Fig. 1A). Histological analysis revealed that male *Panx3* KO mice have higher articular cartilage structural (ACS) scores, indicating damage in the superficial zone of the unmineralized cartilage across the whole tibia compared to SED WT mice (9.361; mean rank difference; p = 0.0165) (Fig. 1B). This is evident by the surface erosion and fibrillation on the medial and lateral tibia. With forced treadmill running, *Panx3* KO mice had larger surface erosions and fibrillation across the joint compared to SED WT mice (16.11; p = 0.0003) (Fig. 1B). When analysing the medial compartment alone, there was weak evidence that SED *Panx3* KO mice had higher ACS scores compared to SED WT mice (5.278; p = 0.1590) (Fig. 1C). However, when subjected to treadmill running, *Panx3* KO mice had larger surface defects compared to SED WT mice (14.78; p = 0.0005) (Fig. 1C). In the lateral compartment, SED *Panx3* KO mice develop superficial cartilage erosion and fibrillation compared to SED WT mice (8.972; p = 0.0171), and strong evidence suggesting differences between FEX *Panx3* KO and SED WT mice (13.60; p = 0.00015) (Fig. 1D). Taken together, this suggests the deletion of *Panx3*, in combination with forced treadmill running, causes greater surface cartilage damage compared to SED WT mice.

**Fig. 1.**
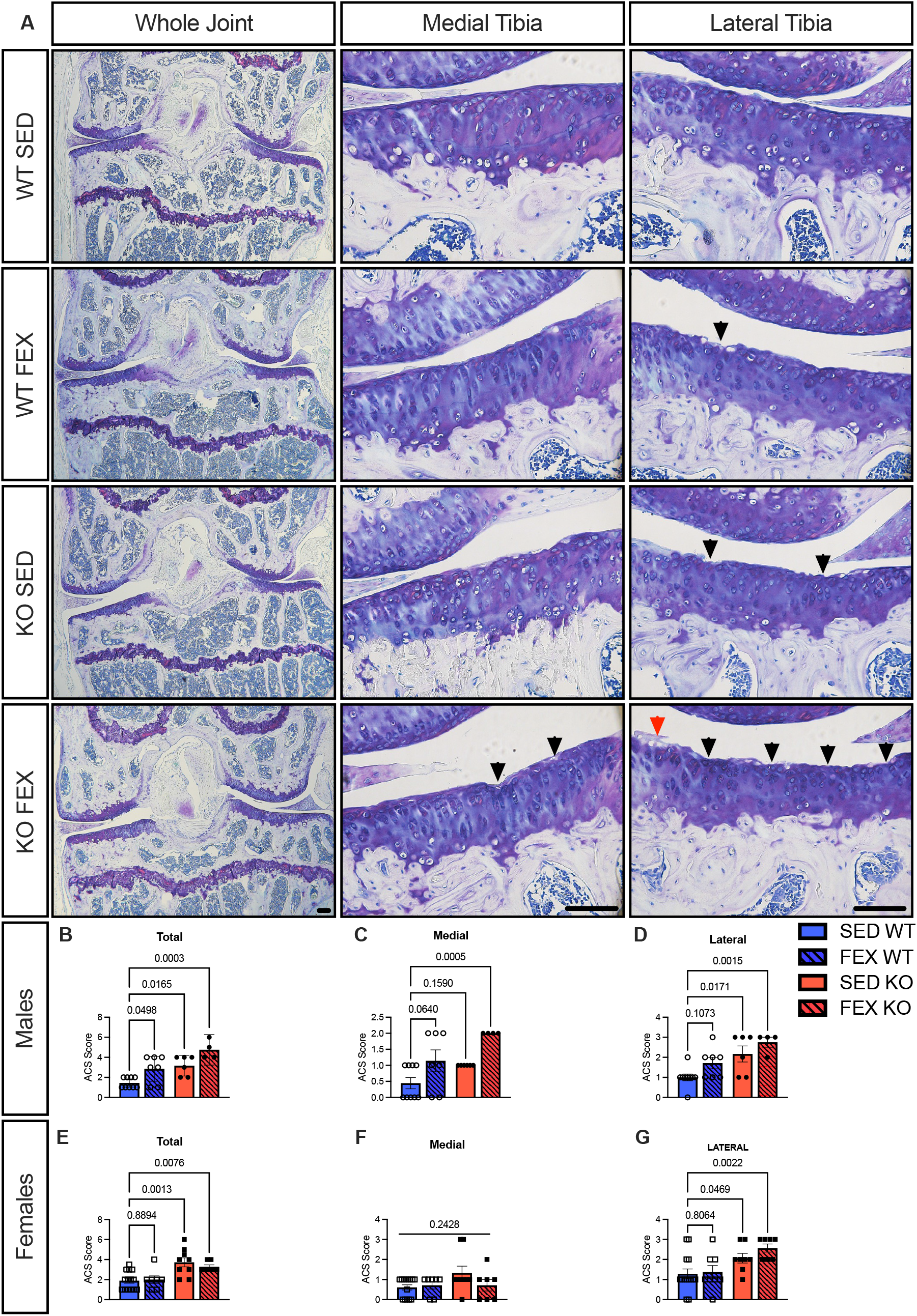
The tibial cartilage of *Panx3* KO mice shows features of OA that are compounded by forced treadmill running. Representative Tolidine Blue staining of knee joints from male and female wildtype (WT) and *Panx3* knockout (KO) mice under sedentary (SED) and forced treadmill running (FEX) conditions, as indicated (**A**). Images on the left show 4x magnification of whole joints in the frontal plane, 20x magnification of tibial cartilage in the medial (middle panel) and lateral (right panel) surfaces. Black arrows indicate cartilage defects, while red arrows indication loss of staining. Scale bars, 100µm. (**B – D**) Total (B), medial (C), and lateral (D) tibial articular cartilage structure (ACS) score of male mice for WT SED (N = 8), WT FEX (N = 7), KO SED (N = 6), and KO FEX (N = 4), as indicated. (**E – G**) Total (E), medial (F), and lateral (G) tibial articular cartilage structure (ACS) score of female mice for WT SED (N = 14), WT FEX (N = 8), KO SED (N = 9), and KO FEX (N = 7), as indicated. For statistical comparisons among the groups, a Kruskal-Wallis test was performed. All data are shown as means ± CI.

For female mice, across the whole tibial joint, statistical and scientific inference would suggest forced treadmill running did not influence ACS scores in either genotype (Fig.1 E-G). There was moderate to strong evidence that both SED (14.44; p = 0.0013) and FEX (13.04; p = 0.0076) *Panx3* KO mice had larger surface defects across the joint compared to SED WT mice (Fig. 1E). In the medial compartment, the ANOVA resulted in a p-value = 0.2428, suggesting there were no structural differences among the groups, and thus we did not run a post-hoc comparison (Fig. 1F). In the lateral compartment, there was moderate evidence that SED *Panx3* KO mice have higher ACS scores compared to SED WT mice (9.125; p = 0.0469), while there was moderate to strong evidence that FEX *Panx3* KO mice had higher ACS scores compared to SED WT (14.71; p = 0.0022) (Fig. 1G).

Next, we analysed the tibial cartilage area and thickness in males (Fig. 2B-E) and females (Fig. 2G-J). Using a three-way ANOVA to analyze the effect of genotype, exercise and sex, we found that there was moderate evidence to indicate that the deletion of *Panx3* reduced the medial tibia unmineralized cartilage area [F (1,56) = 6.108, P = 0.0165] and thickness [F (1,56) = 5.493, p = 0.0227] in both males (Fig. 2B&D) and females (Fig. 2G&I). In the mineralized cartilage of the medial tibia, there was moderate evidence [F (1,56) = 5.229, p = 0.0276] suggesting FEX WT mice have greater mineralized cartilage area compared to SED WT mice (0.0077, 0.0148 to 0.000, p = 0.0276) (Fig. 3B). In the lateral tibia, there was weak evidence for differences in unmineralized cartilage area (Fig. 2C), however, there was moderate evidence for a main effect for activity, suggesting forced treadmill running increased thickness [F (1,53) = 7.992, p = 0.0066] in both males (Fig. 2E) and females (Fig. 2J). Lastly, there was no evidence of differences in the lateral mineralized cartilage area (Fig. 2C & H) or thickness (Fig. 2E & J). Collectively, this data indicates that *Panx3* KO mice have less unmineralized cartilage of the medial tibia, regardless of sex, and develop cartilage damage from forced treadmill running compared to SED WT mice.

**Fig. 2.**
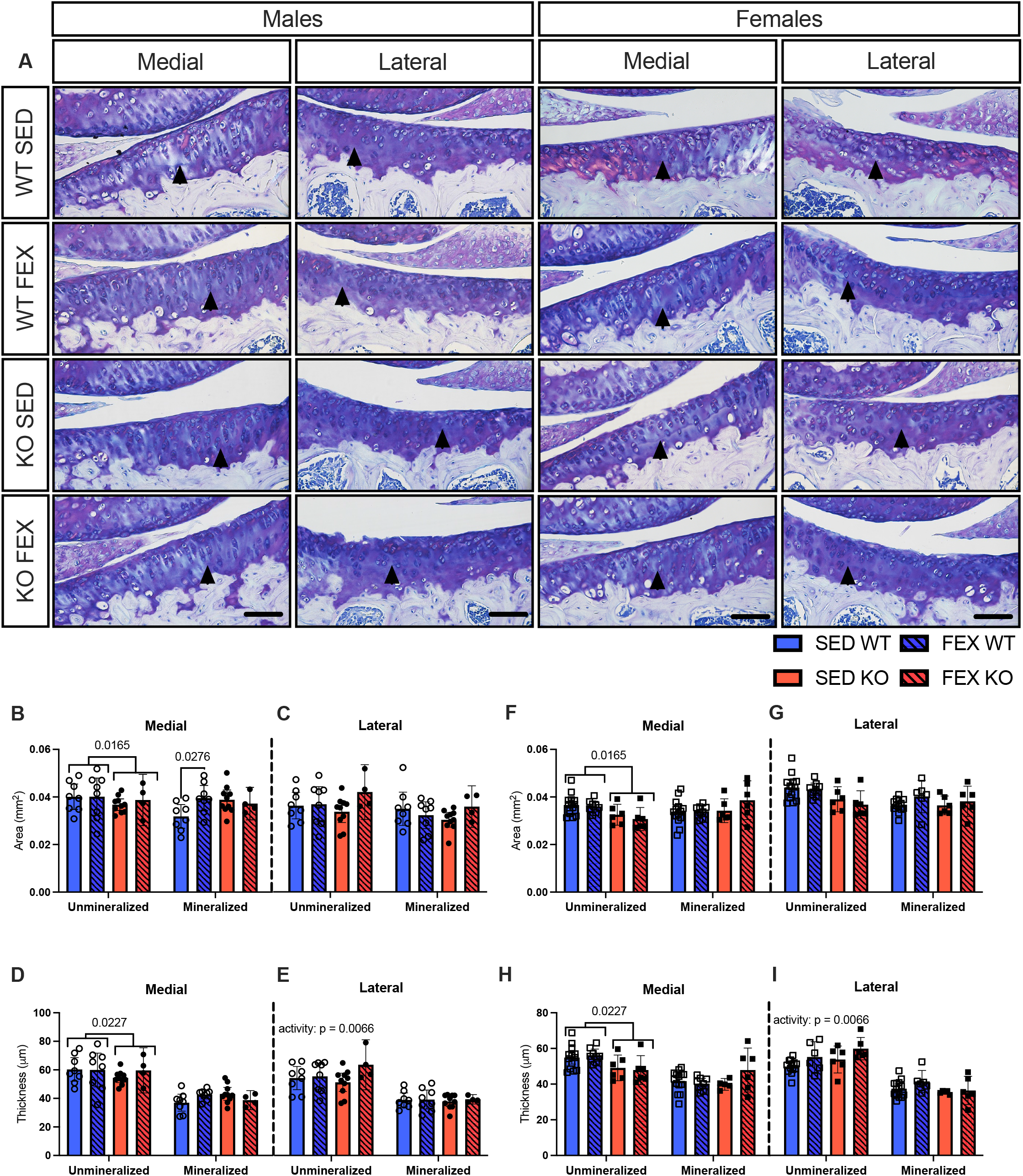
Medial unmineralized cartilage area and thickness of the tibia are reduced in *Panx3* KO mice. Tibial cartilage of the medial and lateral surface was segmented from the underlying subchondral bone and divided into unmineralized and mineralized cartilage for male (**left**) and female (**right**) mice (**A**). Images show 10x magnification of the medial (left) and lateral (right) load bearing regions of the tibial surface. Black arrows indicate tidemark. Medial (**B**) and lateral (**C**) unmineralized area and thickness (**D & E)** for male mice. Medial (**G**) and lateral (**H**) unmineralized area and thickness (**I & K)** for female mice. WT SED (N = 8 males, N = 14 females), WT FEX (N = 8 males, N = 8), KO SED (N = 10 males, N = 6 females), and KO FEX (N = 10 males, N = 6 females), as indicated. For statistical comparisons among the groups, a three-way ANOVA with Sex x Genotype x Activity as factors was used followed by Sidak’s multiple comparisons test. All data are shown as means ± CI.

**Fig. 3.**
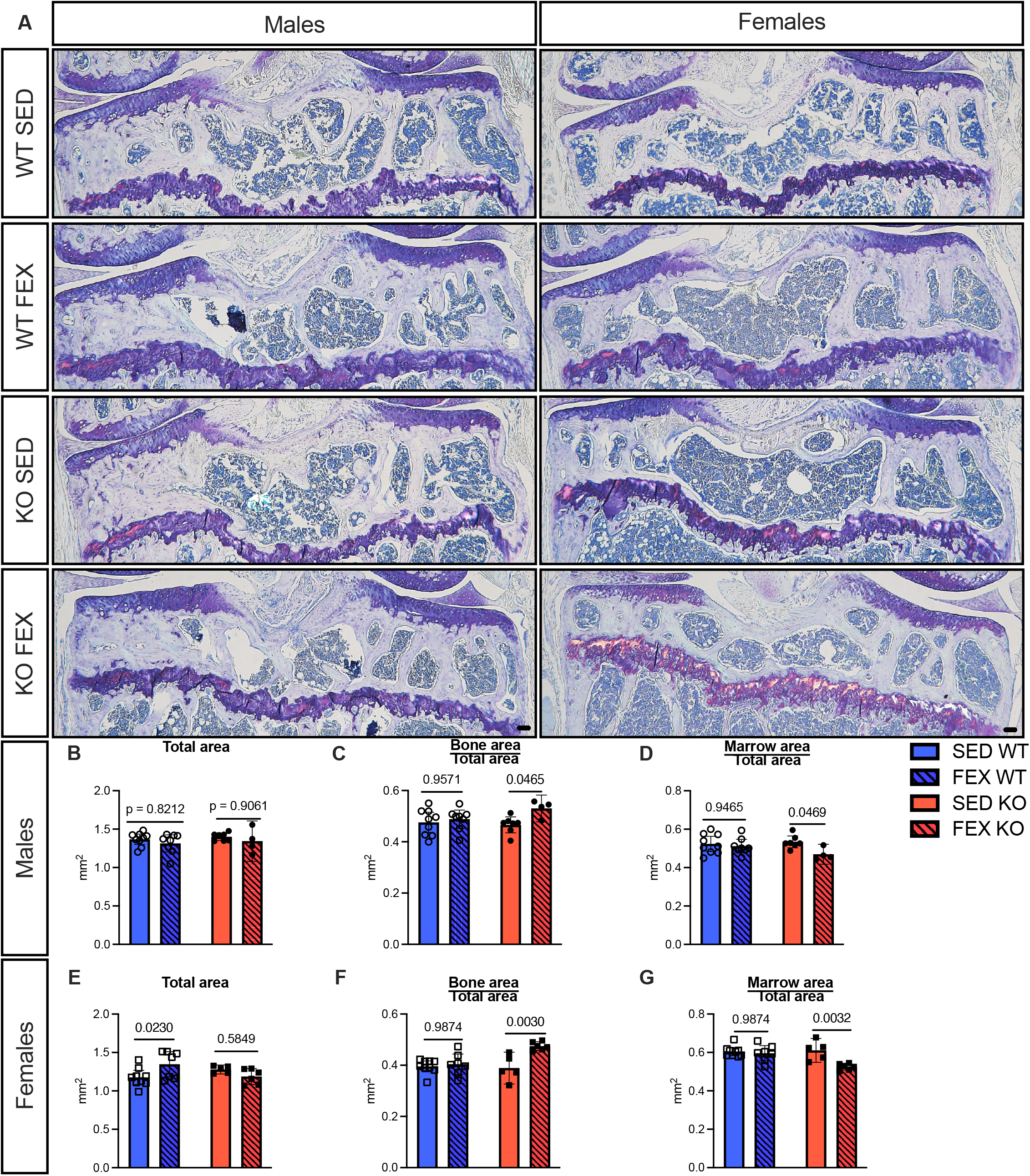
Forced treadmill running results in greater bone of the tibial secondary ossification center in *Panx3* KO mice. Bone and marrow was segmented from the overlaying articular cartilage and the underlying growth plate of the proximal tibia. Representative Tolidine Blue staining from male (left panel) and female (right panel) wildtype (WT) and *Panx3* knockout (KO) mice under sedentary (SED) and forced treadmill running (FEX) conditions, as indicated (**A**). Images show 4x magnification in the frontal plane, Scale bars, 100µm. (**B-D**) Total area (**B**), bone area corrected for total area (**C**), and marrow area corrected for total area (**D**) of male WT and *Panx3* KO mice. WT SED (N = 9), WT FEX (N = 8), KO SED (N = 7), and KO FEX (N = 4), as indicated. (**E-G**) Total area (**E**), bone area corrected for total area (**F**), and marrow area corrected for total area (**G**) of female WT and *Panx3* KO mice. WT SED (N = 9), WT FEX (N = 7), KO SED (N = 5), and KO FEX (N = 6), as indicated. For statistical comparisons among the groups, a three-way ANOVA with Sex x Genotype x Activity as factors followed by Sidak’s multiple comparisons test. All data are shown as means ± CI.

### *Panx3* KO mice have greater bone area in the proximal tibia after forced treadmill running compared to SED *Panx3* KO mice

It has been established that subchondral bone changes of the tibia occur early on in OA [21], and PANX3 is expressed in bone cells and regulates osteoblast differentiation and bone modelling [22]. However, there is no published research investigating the role of *Panx3* in bone remodelling, such as during exercise. Considering that the *Panx3* KO mice develop signs of cartilage erosion, which is exacerbated with treadmill running, we next wanted to determine whether the secondary ossification site in the tibia showed any pathological changes (Fig. 3A). We performed histomorphometry analysis of the proximal tibial secondary ossification site, which involved segmenting the bone from the marrow within the region of interest as explained in the methods section. For male mice, there was no evidence that forced treadmill running altered the total area of interest for either WT (p = 0.8212) or *Panx3* KO (p = 0.5849) mice (Fig. 4B). There was strong evidence for a two-way interaction of genotype x activity [F (1, 47) = 8.660, P = 0.0050] for bone area, resulting in *Panx3* KO mice having increased bone area with forced treadmill running (0.06441; 0.1281, 0.00072); p = 0.0465), while there was no change in WT mice (p = 0.9571) (Fig. 3C). Additionally, there was strong evidence for a two-way interaction of genotype x activity [F (1, 47) = 8.696, P = 0.0050] for marrow area, resulting in moderate evidence that FEX *Panx3* KO mice had less marrow area compared to SED *Panx3* KO mice (0.06442; 0.0006, 0.1282; p = 0.0469), while there was no evidence this change happened in WT mice (p = 0.9571) (Fig. 3D).

**Fig. 4.**
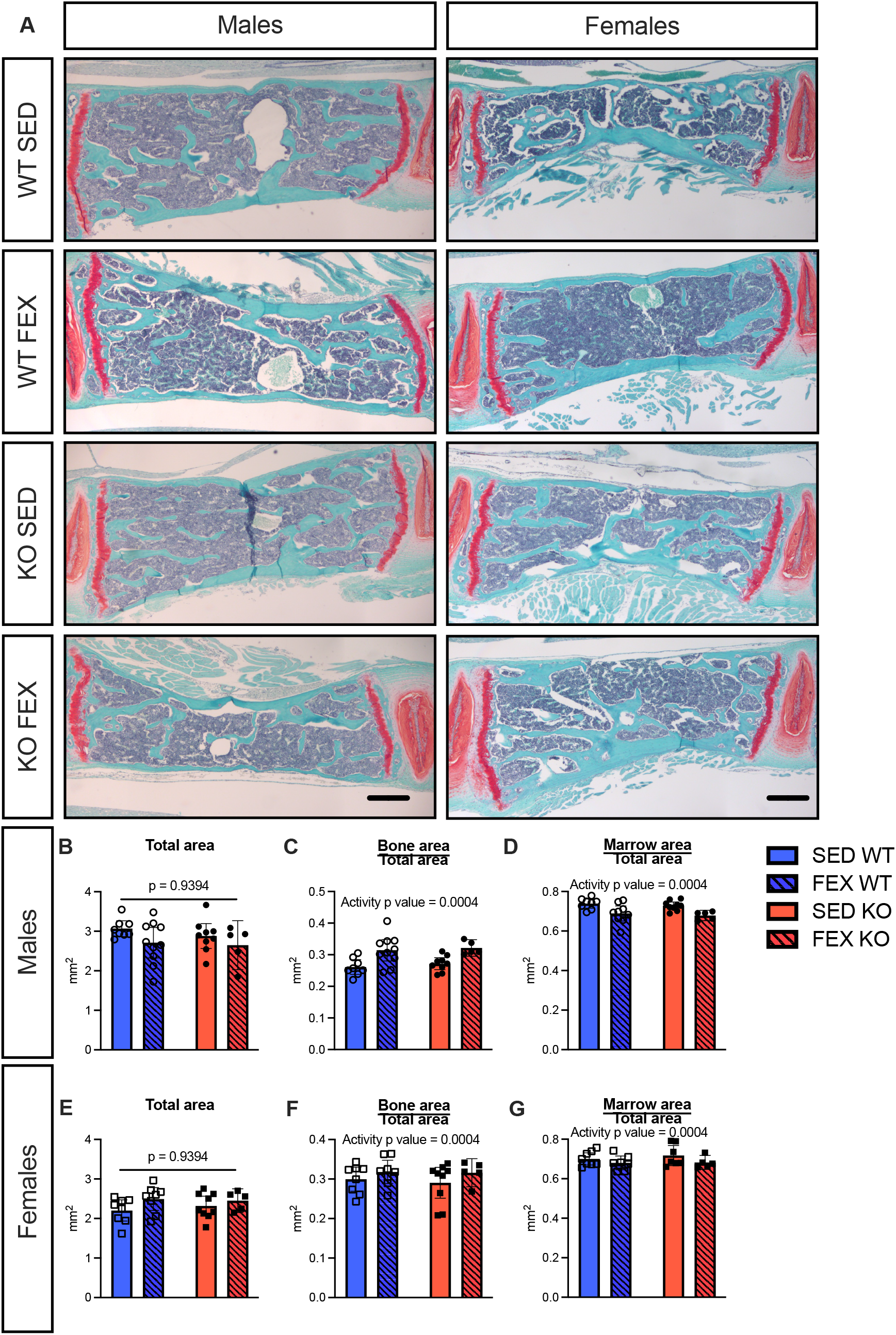
WT and *Panx3* KO mice L5 vertebra respond similarly to forced treadmill running. L5 vertebral bone and marrow was segmented from each other for analysis. Representative Safranin O (SAF O) and Fast Green (FG) staining of the L5 vertebra from male (left panel) and female (right panel) wildtype (WT) and *Panx3* knockout (KO) mice under sedentary (SED) and forced exercise (FEX) conditions, as indicated (**A**). Images show 4x magnification in the frontal plane, Scale bars, 100µm. (**B-D**) Total area (**B**), bone area corrected for total area (**C**), and marrow area corrected for total area (**D**) of male WT and *Panx3* KO mice. WT SED (N = 8), WT FEX (N = 9), KO SED (N = 9), and KO FEX (N = 5), as indicated. (**E-G**) Total area (**E**), bone area corrected for total area (**F**), and marrow area corrected for total area (**G**) of female WT and *Panx3* KO mice. WT SED (N = 8), WT FEX (N = 8), KO SED (N = 9), and KO FEX (N = 5), as indicated. For statistical comparisons among the groups, a three-way ANOVA with Sex x Genotype x Activity as factors. All data are shown as means ± CI.

In female mice, there was moderate evidence that the total secondary ossification site area of the proximal tibia of the WT females increased with forced treadmill running (0.1707, 0.3237, 0.01774; P = 0.0230) (Fig. 4E). Like the male mice, strong evidence suggested that female *Panx3* KO mice had higher bone area with forced treadmill running (0.08566; 0.1472, 0.02413; p = 0.0030) while there was no evidence for this in the WT mice (p = 0.9674) (Fig. 3F). Similarly, there was strong evidence that *Panx3* KO mice have less marrow area with forced treadmill running (0.08538; 0.02376, 0.1470; p = 0.0032) (Fig. 3G). These data suggest that *Panx3* KO mice, regardless of sex, have an altered response of the proximal tibia secondary ossification site to forced treadmill running that may contribute to the development of OA.

### Both WT and *Panx3* KO mice have greater bone area of the L5 vertebral with forced treadmill running compared to their SED counterparts

We next examined the effects of *Panx3* deletion and forced treadmill running on the axial skeleton in this model. We analysed the L5 vertebra as we had done for the tibial secondary ossification site (Fig. 5). There was no evidence that genotype or activity influenced the total area of the L5 vertebrae (Fig. 4B&E). However, there was strong evidence that activity increased bone area (F (1,54) = 14.22, p = 0.0004) (Fig. 4C&F) and decreased marrow area (F (1,54) = 14.58, p = 0.0004) (Fig. 4D&G) in males and females of both genotypes. This would suggest that the forced treadmill running is providing enough stress to induce adaptation in the axial skeleton that is similar between genotypes.

**Fig. 5.**
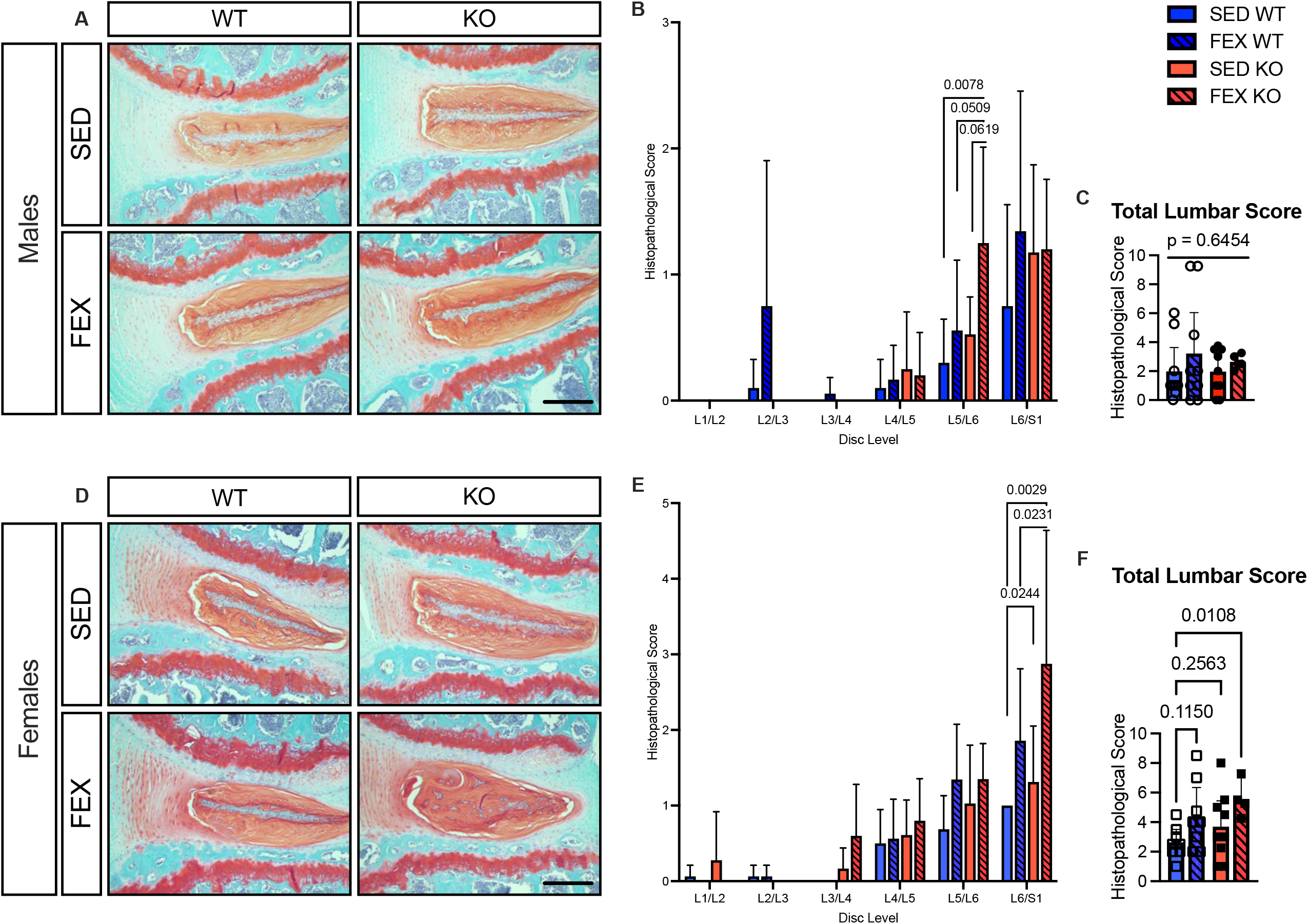
Forced treadmill running *Panx3* KO mice have accelerated IVD degeneration. L5/L6 intervertebral disc (IVD) stained in Safranin O (SAF O) and Fast Green (FG) staining from males (**A**). Images show 10x magnification, scale bars, 100µm. IVD scores separated by disc height (**B**). L5/L6 IVD histopathological score for male mice (**C**). WT SED (N = 10), WT FEX (N = 9), KO SED (N = 10), and KO FEX (N = 5), as indicated. (**D-F**) Female L6/S1 IVD stained in SAF O and FG from female mice (**D**). IVD scores separated by disc height (**E**). L6/S1 IVD histopathological score for female mice (**F**). WT SED (N = 8), WT FEX (N = 8), KO SED (N = 9), and KO FEX (N = 5), as indicated. For statistical comparisons among the groups, a Kruskal-Wallis test was performed. All data are shown as means ± CI.

### Forced treadmill running caused accelerated IVD degeneration in *Panx3* KO mice

We previously elucidated the potential role of PANX3 in maintaining disc homeostasis in both injury and aging models of IVD degeneration [10]. To better understand the role of PANX3 in mediating the response to mechanical load associated with forced exercise, we performed histopathological scoring to assess features of disc degeneration within each compartment of the IVD across the lumbar spine (Fig. 5). While no overt signs of degeneration were detected in the upper lumbar spine, degenerative changes were observed in the lower lumbar regions in both male and female *Panx3* KO mice following forced treadmill running (Fig. 5B&E). In males, there was moderate to strong evidence that under forced treadmill running conditions, *Panx3* KO mice had increased degeneration at the L5/L6 IVD compared to SED WT mice (13.60, p = 0.0078) (Fig. 5B). The increase in histopathological scores were primarily driven by degenerative changes in the annulus fibrosus and interzone, including increased accumulation of hypertrophic cells as well as widening and reversal of the lamellar structure (Fig. 5A). When histopathological scores were combined to assess the entire lumbar spine, there was weak evidence for differences in scores among the groups (p = 0.6454) (Fig. 5C). In females, loss of *Panx3* was likewise associated with IVD degeneration in the lower lumbar spine following forced treadmill running (Fig. 5D). Specifically, moderate to strong evidence suggests FEX *Panx3* KO mice have increased IVD degeneration at L6/S1 compared to FEX WT (9.889, p = 0.0231), and SED WT mice (13.50, p = 0.0029) (Fig. 5E). Additionally, there was moderate evidence that SED *Panx3* KO mice had increased degeneration compared to SED WT mice (8.438, p = 0.0244) (Fig. 5E). When histopathological scores were combined to assess the entire lumbar spine, there was moderate evidence that FEX *Panx3* KO mice had increased degeneration compared to SED WT mice (13.25, p = 0.0108) (Fig. 5F). Following forced exercise, IVDs from *Panx3* KO mice showed hallmark degenerative changes in the annulus fibrosus and interzone, such as hypertrophic cells and widening and reversal of the lamellar structure compared to SED WT mice. This data suggests that forced treadmill running in *Panx3* KO mice causes accelerated IVD degeneration in both male and female mice.

## DISCUSSION

Here, we characterized the effects of forced treadmill running on knee joints and lumbar spines of male and female *Panx3* KO mice. Both male and female *Panx3* KO mice have thinner unmineralized cartilage of the medial tibial compared to WT mice. With the addition of forced treadmill running, *Panx3* KO mice showed large surface defects of the unmineralized cartilage and alterations of the secondary ossification center in the tibia. In addition to these changes to the knee, the forced treadmill running stressed the axial skeleton, as evidenced by the greater bone area in the L5 compared to sedentary mice for both WT and *Panx3* KO mice. This stress, in both male and female *Panx3* KO mice, was associated with accelerated IVD degeneration at L5/L6 and L6/S1 disc levels, respectively, while there was weak evidence for this effect in WT mice. Collectively, this data indicates that the loss of *Panx3* in mice, in combination with 6 weeks of forced treadmill running, induces pathological changes to joint tissues.

We have previously shown that male *Panx3* KO mice are protected from a traumatic model of OA [8], but demonstrate worse OA during aging [9]. In our previous study of DMM surgery in *Panx3* KO mice, we only used male animals, and therefore the results may not be representative of females. Additionally, considering the severity of OA in DMM models, we did not observe the subtle differences in cartilage structure that may have been present in the non-surgical knees of the *Panx3* KO mice. For the present study, we used the 12-point articular cartilage structure scoring system which seems to be more sensitive to the subtle differences that may be present in OA models such as forced treadmill running [17]. It seems that while *Panx3* KO mice are protected from post-traumatic OA, the present data, and our aging study [9] would suggest these mice develop accelerated spontaneous OA and are sensitive to the mechanical stress of forced treadmill running.

Forced treadmill running offers an interesting intervention to study the effects of genetic factors on OA and IDD, as there is a complex relationship between mechanical use and the development of pain and joint radiographic pathology. This relationship seems to be U-shaped in nature in humans, as OA is more prevalent in sedentary populations [23], and there is an increased risk associated with frequent high volumes of loading, as seen in the workplace [24]. The degree to which one is susceptible to mechanical “overuse” could be influenced by genetic differences. A limited number of genetic studies have used forced treadmill running interventions in OA [14, 25-27], as most studies use more traumatic models [28]. Bomer et al., however, determined that forced treadmill running caused differential expression of multiple OA associated genes in cartilage, validating its use as a model of OA [14]. Subsequently, they determined that mice lacking the *DIO2* gene (previously identified as a susceptible locus for OA) were protected from treadmill running induced OA. This suggests genetic factors may in fact influences one’s susceptibility to exercise induced OA. Matsuzaki et al., performed aging, DMM and treadmill running in a Col2Cre-FoxO KO mouse model to determine the importance of this transcription factor in various OA contexts [26], while Rellman et al. showed that in protein disulfide isomerase ERp57 KO, to induce chondrocyte ER stress, mice are susceptible to age but not forced treadmill induced OA [27]. Studies like these are important as they highlight the molecular differences between various factors that influence OA progression. Considering studies have shown that exercise can induce knee OA [29-31], and paradoxically, studies also show exercise reduces OA development [25, 32] in mice, it would be imperative to elucidate the genetic factors, and also the exercise parameters that elicit these contrasting results. Here we implemented a modified forced treadmill running protocol previously reported to induce knee OA in mice [14]. Surprisingly, we observed weak to moderate evidence that this protocol induced OA in our WT mice. Additionally, in female mice, our model suggests forced treadmill running had little to no effect on knee cartilage structure within each genotype. We are unaware of any previous studies investigating the effect of forced treadmill running on female C57BL6 mouse cartilage to compare these findings. Considering that we saw the greatest development of OA in the *Panx3* KO mice that were forced to run, these data may suggest an interaction or additive effect between activity and genotype. However, the nature of this data does not allow for the performance of statistical tests to determine such relationships.

With regards to IVDs, we have previously examined the influence of *Panx3* deletion on IDD in both aging and IVD puncture (traumatic) models [10]. While there were no obvious differences in pathological IVD changes between *Panx3* KO and WT mice during aging, following IVD puncture, AF tissue architecture appeared better preserved in IVDs from *Panx3* KO mice compared to WT mice. This would suggest that the absence of *Panx3* in traumatic injuries is beneficial. Interestingly, IVDs adjacent to the site of puncture, which experience increased mechanical stress, showed signs of accelerated nucleus pulposus degeneration [10]. These data suggest that *Panx3* KO mice are prone to accumulating damage of the lumbar IVD when mechanically stressed. Considering we saw increased IVD pathology in our forced treadmill running KO mice, this supports this previous finding, and suggests these mice are sensitive to mechanical stress. Interestingly, Belonogova et al. found that *PANX3* rare polymorphic non-coding variants in humans are strongly associated with back pain, which suggests PANX3 may be involved in human IDD associated pain [11]. Considering our findings using our *Panx3* KO mouse model, and the findings of Belonogova et al. in humans [11], further investigation of PANX3 in IVD health and disease is warranted.

To our knowledge, there are no forced treadmill running studies investigating its effects on IVD health in mice, as all the rodent studies are in rats [33-35]. Rat models of forced treadmill running have shown to be anabolic for IVDs [34], and dynamic compressive forces to IVDs in an *in vitro* model stimulated an anabolic response by increasing gene expression for types I and II collagen and aggrecan [36]. Additionally, other researchers have shown that the effect of hydrostatic pressure on IVD response is dose dependent, with higher forces eliciting a catabolic response [37]. These data suggest dynamic, cyclical, loading (such as during running) may be anabolic for IVDs, which is supported by cross-sectional studies in humans [38, 39]. In our model, *Panx3* KO mice forced to run develop histological features of IDD while there was weak evidence for this in WT mice. Like the knee data, this would suggest that the *Panx3* KO mice are sensitive to the mechanical stress of the running. What molecular mechanism is mediating this potentially pathological response is to be determined.

Unfortunately, our model had low n values for multiple groups which increased the range of our confidence intervals. Considering the effect of the forced treadmill running on OA and IDD measured in either genotype is small (if any in the female WT mice), our experiment provides weak statistical evidence to determine, with a high degree of confidence, the precise effect of forced treadmill running within each genotype. We had lost 2 male FEX *Panx3* KO samples during processing which ended up being the rate limiting group for the statistical analysis. However, considering we analysed multiple tissues and sites at the knee joint, and IVDs, in both males and females, which all provided similar relationships among the groups, our data suggests that the combination of deleting *Panx3* and forced treadmill running worsened OA and IDD measures. In fact, if we were to pool males and females within their respective groups, this further strengthens the statistical inference that the genotype in combination with forced treadmill running exacerbates joint pathology (data not shown). Additionally, the OA and IDD differences we observed in our analysis are relatively mild compared to phenotypes observed in other genetic and traumatic injury induced models. This could be due, in part, to the parameters of the treadmill running protocol. Nevertheless, the clinical impacts of these findings are therefore uncertain, and further studies investigating permutations of deleting *Panx3* under spontaneous models are needed. It is possible that older mice may be more susceptible to the effects of forced treadmill running, which may exacerbate the differences between genotypes.

A strength of our model was the consideration of both male and female mice. While WT and *Panx3* KO mice were not littermates, littermate siblings within each genotype were divided between activity groups. We believe this is an important control to determine the effect of the forced treadmill running within each genotype. This study was not powered a priori to detect statistically significant changes in OA or IVD measures. Thus, these statistical analyses should be interpreted with caution. Lastly, while our model determined that *Panx3* KO mice forced to run had superficial cartilage lesions in the tibia of the knee and IVD histopathology, it is known that structured exercise is effective at reducing OA pain and improve function, which we did not measure. Therefore, function and pain-related changes due to the structural differences are unknown in these mice. Additionally, we did not measure voluntary activity of the mice while in the cages, and it is possible that there were differences in activity between genotypes that was not documented.

## Conclusion

These findings indicate that PANX3 is important for the response of musculoskeletal tissues to the stress of forced treadmill running in mice. This work is in line with our previous findings on the role of PANX3 in OA and the effect of various interventions on knee and IVD tissue response, that could be further explored as new targets for disease interventions.

## Abbreviations

OA: Osteoarthritis
IDD: Intervertebral disc disease
IVD: Intervertebral disc
Panx3: Pannexin 3
KO: Knockout
WT: Wildtype
SED: Sedentary
FEX: Forced exercise
DMM: Destabilization of the medial meniscus

## Notes

### Competing Interest Statement

The authors have declared no competing interest.

## REFERENCES

1. Cui, A., et al., Global, regional prevalence, incidence and risk factors of knee osteoarthritis in population-based studies. EClinicalMedicine, 2020. 29-30: p. 100587.

2. Wu, A., et al., Global low back pain prevalence and years lived with disability from 1990 to 2017: estimates from the Global Burden of Disease Study 2017. Ann Transl Med, 2020. 8(6): p. 299.

3. Freidin, M.B., et al., Insight into the genetic architecture of back pain and its risk factors from a study of 509,000 individuals. Pain, 2019. 160(6): p. 1361–1373.

4. Alzahrani, H., et al., The association between physical activity and low back pain: a systematic review and meta-analysis of observational studies. Scientific Reports, 2019. 9(1): p. 8244.

5. Ishikawa, M., et al., Pannexin 3 ER Ca(2+) channel gating is regulated by phosphorylation at the Serine 68 residue in osteoblast differentiation. Sci Rep, 2019. 9(1): p. 18759.

6. Iwamoto, T., et al., Pannexin 3 regulates intracellular ATP/cAMP levels and promotes chondrocyte differentiation. J Biol Chem, 2010. 285(24): p. 18948–58.

7. Veras, M.A., et al., Transcriptional profiling of the murine intervertebral disc and age-associated changes in the nucleus pulposus. Connect Tissue Res, 2020. 61(1): p. 63–81.

8. Moon, P.M., et al., Deletion of Panx3 Prevents the Development of Surgically Induced Osteoarthritis. J Mol Med (Berl), 2015. 93(8): p. 845–56.

9. Moon, P.M., et al., Global Deletion of Pannexin 3 Resulting in Accelerated Development of Aging-Induced Osteoarthritis in Mice. Arthritis & Rheumatology, 2021. 73(7): p. 1178–1188.

10. Serjeant, M., et al., The Role of Panx3 in Age-Associated and Injury-Induced Intervertebral Disc Degeneration. Int J Mol Sci, 2021. 22(3).

11. Belonogova, N.M., et al., Noncoding rare variants in PANX3 are associated with chronic back pain. Pain, 2022.

12. Bond, S.R., et al., Pannexin 3 is a novel target for Runx2, expressed by osteoblasts and mature growth plate chondrocytes. J Bone Miner Res, 2011. 26(12): p. 2911–22.

13. van der Kraan, P.M. and W.B. van den Berg, Chondrocyte hypertrophy and osteoarthritis: role in initiation and progression of cartilage degeneration? Osteoarthritis Cartilage, 2012. 20(3): p. 223–32.

14. Bomer, N., et al., The effect of forced exercise on knee joints in Dio2(-/-) mice: type II iodothyronine deiodinase-deficient mice are less prone to develop OA-like cartilage damage upon excessive mechanical stress. Ann Rheum Dis, 2016. 75(3): p. 571–7.

15. Wakefield, C.B., et al., Pannexin 3 deletion reduces fat accumulation and inflammation in a sex-specific manner. International Journal of Obesity, 2022. 46(4): p. 726–738.

16. Abitbol, J.M., et al., Double deletion of Panx1 and Panx3 affects skin and bone but not hearing. J Mol Med (Berl), 2019. 97(5): p. 723–736.

17. McNulty, M.A., et al., A Comprehensive Histological Assessment of Osteoarthritis Lesions in Mice. Cartilage, 2011. 2(4): p. 354–63.

18. McCann, M.R., et al., Repeated exposure to high-frequency low-amplitude vibration induces degeneration of murine intervertebral discs and knee joints. Arthritis Rheumatol, 2015. 67(8): p. 2164–75.

19. Tam, V., et al., Histological and reference system for the analysis of mouse intervertebral disc. J Orthop Res, 2018. 36(1): p. 233–243.

20. Wasserstein, R.L., A.L. Schirm, and N.A. Lazar, Moving to a World Beyond “p < 0.05”. American Statistician, 2019. 73: p. 1–19.

21. Fang, H., et al., Early Changes of Articular Cartilage and Subchondral Bone in The DMM Mouse Model of Osteoarthritis. Scientific Reports, 2018. 8(1): p. 2855.

22. Plotkin, L.I., et al., Connexins and Pannexins in Bone and Skeletal Muscle. Curr Osteoporos Rep, 2017. 15(4): p. 326–334.

23. Yukalang, N., et al., Association Between Physical Activity and Osteoarthritis of Knee with Quality of Life in Community-Dwelling Older Adults. Stud Health Technol Inform, 2021. 285: p. 265–270.

24. Parsons, C.M., et al., Predominant lifetime occupation and associations with painful and structural knee osteoarthritis: An international participant-level cohort collaboration. Osteoarthr Cartil Open, 2020. 2(4): p. 100085.

25. Pate, K.M., et al., The beneficial effects of exercise on cartilage are lost in mice with reduced levels of ECSOD in tissues. J Appl Physiol (1985), 2015. 118(6): p. 760–7.

26. Matsuzaki, T., et al., FoxO transcription factors modulate autophagy and proteoglycan 4 in cartilage homeostasis and osteoarthritis. Sci Transl Med, 2018. 10(428).

27. Rellmann, Y., et al., ER Stress in ERp57 Knockout Knee Joint Chondrocytes Induces Osteoarthritic Cartilage Degradation and Osteophyte Formation. Int J Mol Sci, 2021. 23(1).

28. Fang, H. and F. Beier, Mouse models of osteoarthritis: modelling risk factors and assessing outcomes. Nature Reviews Rheumatology, 2014. 10(7): p. 413–421.

29. Zhou, X., et al., Moderate-intensity treadmill running relieves motion-induced post-traumatic osteoarthritis mice by up-regulating the expression of lncRNA H19. BioMedical Engineering OnLine, 2021. 20(1): p. 111.

30. Lapveteläinen, T., et al., Lifelong moderate running training increases the incidence and severity of osteoarthritis in the knee joint of C57BL mice. Anat Rec, 1995. 242(2): p. 159–65.

31. Yamagishi, K., et al., Activation of the renin-angiotensin system in mice aggravates mechanical loading-induced knee osteoarthritis. Eur J Histochem, 2018. 62(3).

32. Huesa, C., et al., Moderate exercise protects against joint disease in a murine model of osteoarthritis. Front Physiol, 2022. 13: p. 1065278.

33. Luan, S., et al., Running exercise alleviates pain and promotes cell proliferation in a rat model of intervertebral disc degeneration. Int J Mol Sci, 2015. 16(1): p. 2130–44.

34. Brisby, H., et al., The effect of running exercise on intervertebral disc extracellular matrix production in a rat model. Spine (Phila Pa 1976), 2010. 35(15): p. 1429–36.

35. Sasaki, N., et al., Physical exercise affects cell proliferation in lumbar intervertebral disc regions in rats. Spine (Phila Pa 1976), 2012. 37(17): p. 1440–7.

36. Wang, D.L., S.D. Jiang, and L.Y. Dai, Biologic response of the intervertebral disc to static and dynamic compression in vitro. Spine (Phila Pa 1976), 2007. 32(23): p. 2521–8.

37. Neidlinger-Wilke, C., et al., Regulation of gene expression in intervertebral disc cells by low and high hydrostatic pressure. Eur Spine J, 2006. 15 Suppl 3(Suppl 3): p. S372–8.

38. Belavý, D.L., et al., Running exercise strengthens the intervertebral disc. Sci Rep, 2017. 7: p. 45975.

39. Mitchell, U.H., et al., Long-term running in middle-aged men and intervertebral disc health, a cross-sectional pilot study. PLoS One, 2020. 15(2): p. e0229457.

